# Perspectives from animal demography on incorporating evolutionary mechanism into plant population dynamics

**DOI:** 10.1101/420869

**Authors:** Maria Paniw

## Abstract

With a growing number of long-term, individual-based data on natural populations available, it has become increasingly evident that environmental change affects populations through complex, simultaneously occurring demographic and evolutionary processes. Analyses of population-level responses to environmental change must therefore integrate demography and evolution into one coherent framework. Integral projection models (IPMs), which can relate genetic and phenotypic traits to demographic and population-level processes, offer a powerful approach for such integration. However, a rather artificial divide exists in how plant and animal population ecologists use IPMs. Here, I argue for the integration of the two sub-disciplines, particularly focusing on how plant ecologists can diversify their toolset to investigate selection pressures and eco-evolutionary dynamics in plant population models. I provide an overview of approaches that have applied IPMs for eco-evolutionary studies and discuss a potential future research agenda for plant population ecologists. Given an impending extinction crisis, a holistic look at the interacting processes mediating population persistence under environmental change is urgently needed.

## Introduction

The environment affects population dynamics through complex effects on demographic rates such as survival and reproduction (Ozgul *et al.*, 2010; Paniw *et al.*, 2018). These ecological processes are, in turn, mediated by differential responses of phenotypes to environmental drivers; and the evolutionary mechanisms underlying such responses can themselves be altered by ecological processes, creating eco-evolutionary feedbacks (Schoener, 2011; Johnston *et al.*, 2013; Lowe *et al.*, 2017). With the realization that evolution can occur rapidly, on the time scale of ecological processes (Schoener, 2011; Ellner, 2013), and that eco-evolutionary feedbacks can strongly affect population responses to environmental change, it has increasingly become evident that a framework is required to flexibly link quantitative traits and population dynamics (Smallegange & Coulson, 2013; Vindenes & Langangen, 2015).

To assess simultaneously occurring, complex feedbacks of ecology (environmental variation and demography) and evolution (changes in static and dynamic traits), population ecologists require (i) individual-level data on traits; (ii) information on transmission, or inheritance, of the traits; (iii) a demographic model that describes demographic rates and transitions as functions of these traits; (iv) a flexible modeling framework that assesses population dynamics from the demographic model and can integrate environmental (e.g., climate, flooding, or disease outbreaks) and intrinsic (e.g., population density) factors (Rees & Ellner, 2016). Integral projection models (IPMs) have emerged as just such a framework (Smallegange & Coulson, 2013; Vindenes & Langangen, 2015). In their basic formulation, these models link continuous distributions of phenotypic traits that change dynamically over time to demographic rates. IPMs can therefore simultaneously project trait and population dynamics (Easterling *et al.*, 2000; Coulson *et al.*, 2010; Smallegange & Coulson, 2013; Merow *et al.*, 2014a). The relationship between vital rates and the continuous trait variable is typically expressed with (generalized) linear models. For example, one can define individual size as the trait and model size at time *t* +1 (*z’*) as a function of size at time *t* (*z*) using a simple regression:

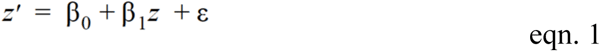

The key flexibility of IPMs lies in the fact that demographic rates can also differ in time and among different ages, stages, or environmental states (Ellner & Rees, 2006; Rees & Ellner, 2009); be made a function of intrinsic population processes such as density feedbacks (Metcalf *et al.*, 2008) or interactions with other species (Adler *et al.*, 2010, 2012); and include static phenotypic and genetic traits such as the genotype of an individual or mass at birth, which do not change through time (see chapter 9 in Ellner *et al.*, 2016; Vindenes & Langangen, 2015).

The demographic rates are then incorporated into a projection kernel *K*(*z′,z*), describing, to follow the example above, the probability of *z*-sized individuals to transition to size *z*’ within one discrete time step. A full *K* consists of two subkernels:

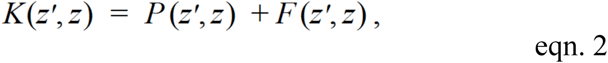

where *P*(*z′,z*) describes the probability of a *z*-sized individual to survive and, conditional on survival, grow to *z*’, and *F*(*z′,z*) describes the probability of a *z*-sized individual to produce offspring with a *z*’ size distribution (Fig. 1; Table 1). As the subkernels *P* and *F* describe a continuous density distribution of *z*, integration is required to obtain *z*’ at *t*+1. To achieve this integration, a midpoint rule is typically applied, which discretizes the kernels into bins over a range of *z* values bounded by the lowest (L) and highest (U) observed values. The kernel *K* is then multiplied by *n*, the number of individuals in the size range [*z, z*+*dz*] to obtain the distribution of *z*’-sized individuals at time *t* +1:

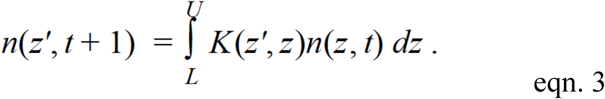

**Table 1.**
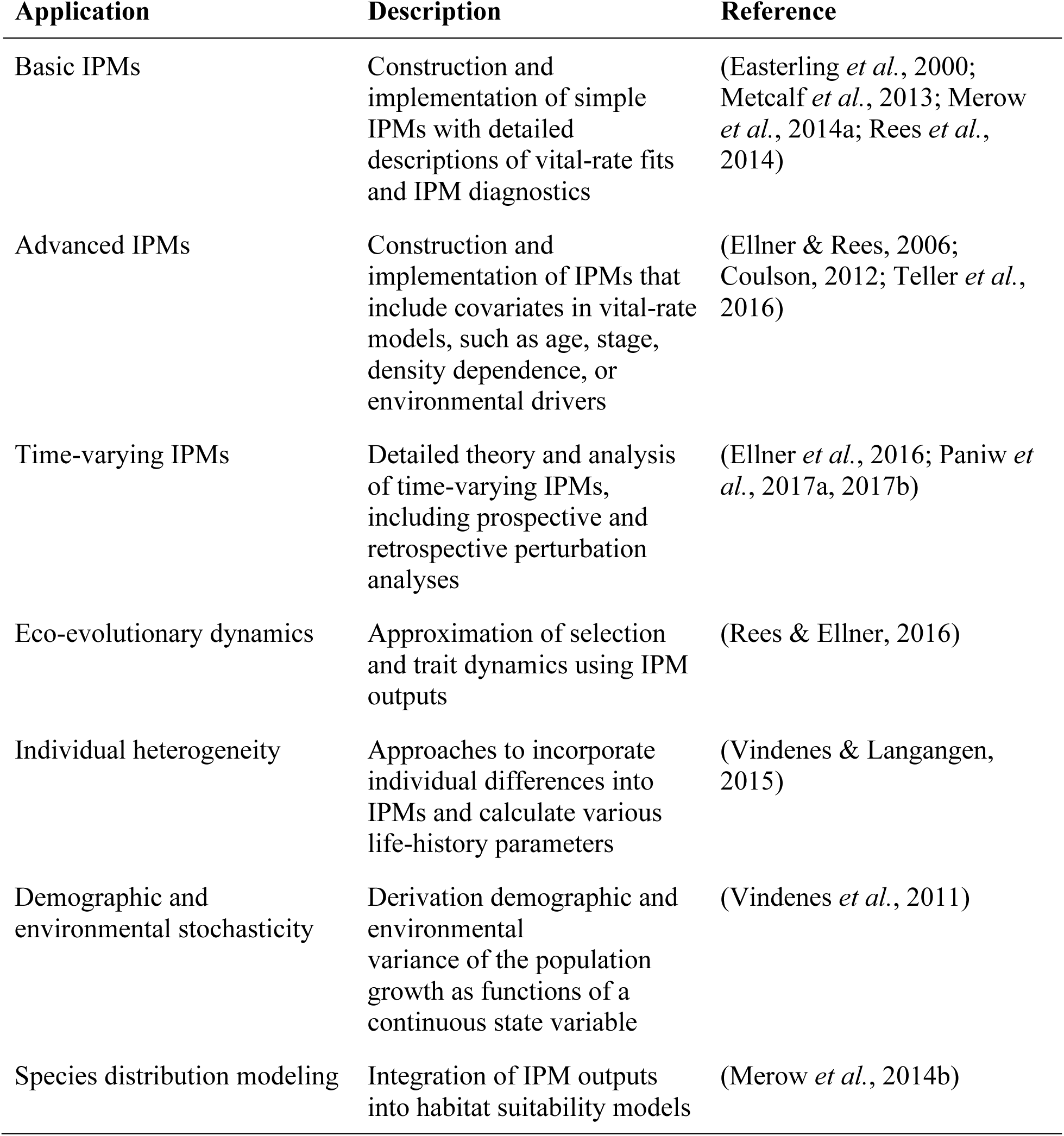
A non-exhaustive summary of applications of integral projection models (IPMs) for which detailed *R* scripts are available.

| Application | Description | Reference |
| --- | --- | --- |
| Basic IPMs | Construction and implementation of simple IPMs with detailed descriptions of vital-rate fits and IPM diagnostics | (Easterling *et al.*, 2000; Metcalf *et al.*, 2013; Merow *et al.*, 2014a; Rees *et al.*, 2014) |
| Advanced IPMs | Construction and implementation of IPMs that include covariates in vital-rate models, such as age, stage, density dependence, or environmental drivers | (Ellner & Rees, 2006; Coulson, 2012; Teller *et al.*, 2016) |
| Time-varying IPMs | Detailed theory and analysis of time-varying IPMs, including prospective and retrospective perturbation analyses | (Ellner *et al.*, 2016; Paniw *et al.*, 2017a, 2017b) |
| Eco-evolutionary dynamics | Approximation of selection and trait dynamics using IPM outputs | (Rees & Ellner, 2016) |
| Individual heterogeneity | Approaches to incorporate individual differences into IPMs and calculate various life-history parameters | (Vindenes & Langangen, 2015) |
| Demographic and environmental stochasticity | Derivation demographic and environmental variance of the population growth as functions of a continuous state variable | (Vindenes *et al.*, 2011) |
| Species distribution modeling | Integration of IPM outputs into habitat suitability models | (Merow *et al.*, 2014b) |

**Figure 1:**
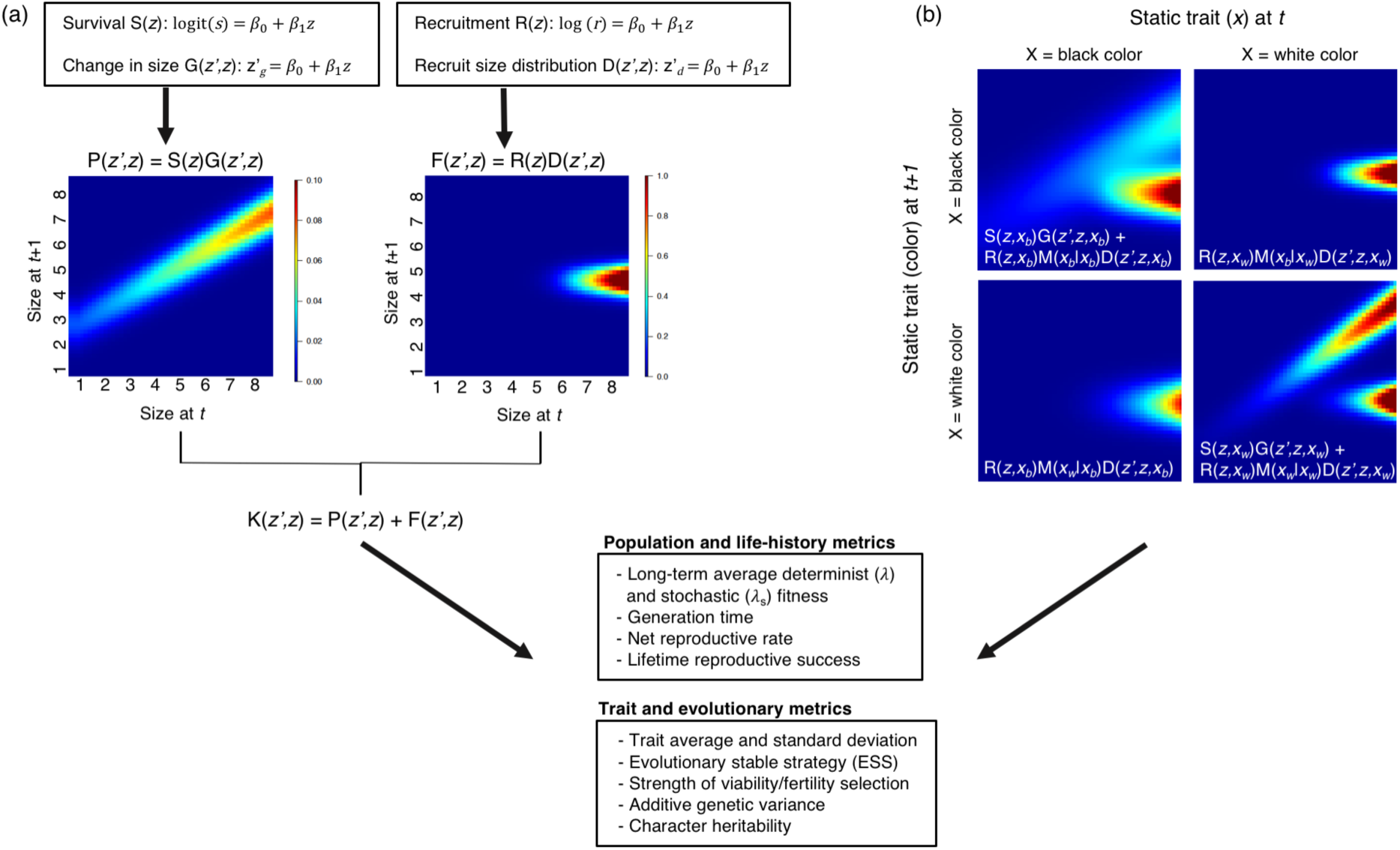
Example of the construction and output of integral projection models (IPMs) for eco-evolutionary dynamics. In a simple IPM (a), demographic rates describing survival, changes in traits (e.g., size), reproduction/recruitment, and offspring development can be assessed as functions of a continuous (phenotypic) trait *z*, via simple (generalized) linear models and integrated into discretized IPM projection kernels, which describe the probability of an individual in the *z*-trait class to transition to the *z*’-trait class. Matrix approximation of the discretized IPMs can be used to calculate population-level metrics, and if inheritance of *z*’-trait offspring by *z*-trait parents is established, evolutionary metrics can also be obtained. In an IPM integrating a static genetic trait *x* (e.g., fur color or size at birth) in addition to a continuous phenotype (b), different kernels can be fit for each genotype, and genetic inheritance *M* can be expressed in terms of genotype e.g., (*x*_*b*_| *x*_*w*_).

Aside from the flexibility of IPMs, their growing popularity is due to the many tutorials on the construction and analysis of these models using the freely available statistical program *R* (R Core Team, 2017; Table 1; see in particular Ellner *et al.*, 2016).

Despite the potential broad applications of IPMs for ecological and global-change research, there has been a rather artificial disconnect in how trait dynamics are studied in animal and plant populations (Smallegange & Coulson, 2013; Ellner *et al.*, 2016). The aim of this article is therefore to bridge this divide by encouraging both sub-disciplines to take a broader perspective in available methodology to advance research linking eco-evolutionary processes to population dynamics. More precisely, I encourage plant ecologists to look more closely to animal population ecology, which offers novel avenues of tackling important aspects of population responses to environmental change.

## Studying the outcome vs. the process of eco-evolutionary dynamics with IPMs

### Evolutionary endpoints

Using structured population models, including IPMs, two main approaches have emerged to investigate eco-evolutionary feedbacks resulting from environmental change: assessing evolutionary outcomes of such change, or evolutionary stable strategies (ESS), and investigating dynamic changes in trait distributions as the environment changes (Fig. 2). In plant populations, eco-evolutionary applications of IPMs have focused strongly on ESS (Geritz *et al.*, 1998; Hendry, 2016; Govaert *et al.*, 2018). The main aim has been to find trait values that maximize long-term fitness and render a given population immune to invasion by mutants, often with respect to reproduction, e.g., the evolution of delayed reproduction and bet hedging (Hesse *et al.*, 2008; Metcalf *et al.*, 2008; Gremer & Venable, 2014; but see Childs *et al.*, 2011 for an example in an animal population). While the assumptions about the genetic basis for such traits are relatively simple (Toprak *et al.*, 2011; Becks *et al.*, 2012), IPMs have allowed researchers to incorporate complex relationships between traits and demographic rates when estimating ESS, including density feedbacks (Ellner *et al.*, 2016). For instance, Metcalf and coauthors (2008) used IPMs to assess life-history decisions, in terms of onset of flowering, in the thistles *Carlina vulgaris* and *Carduus nutans*; and determined accurate predictions of such decisions are possible using long-term empirical, individual-level datasets.

**Figure 2:**
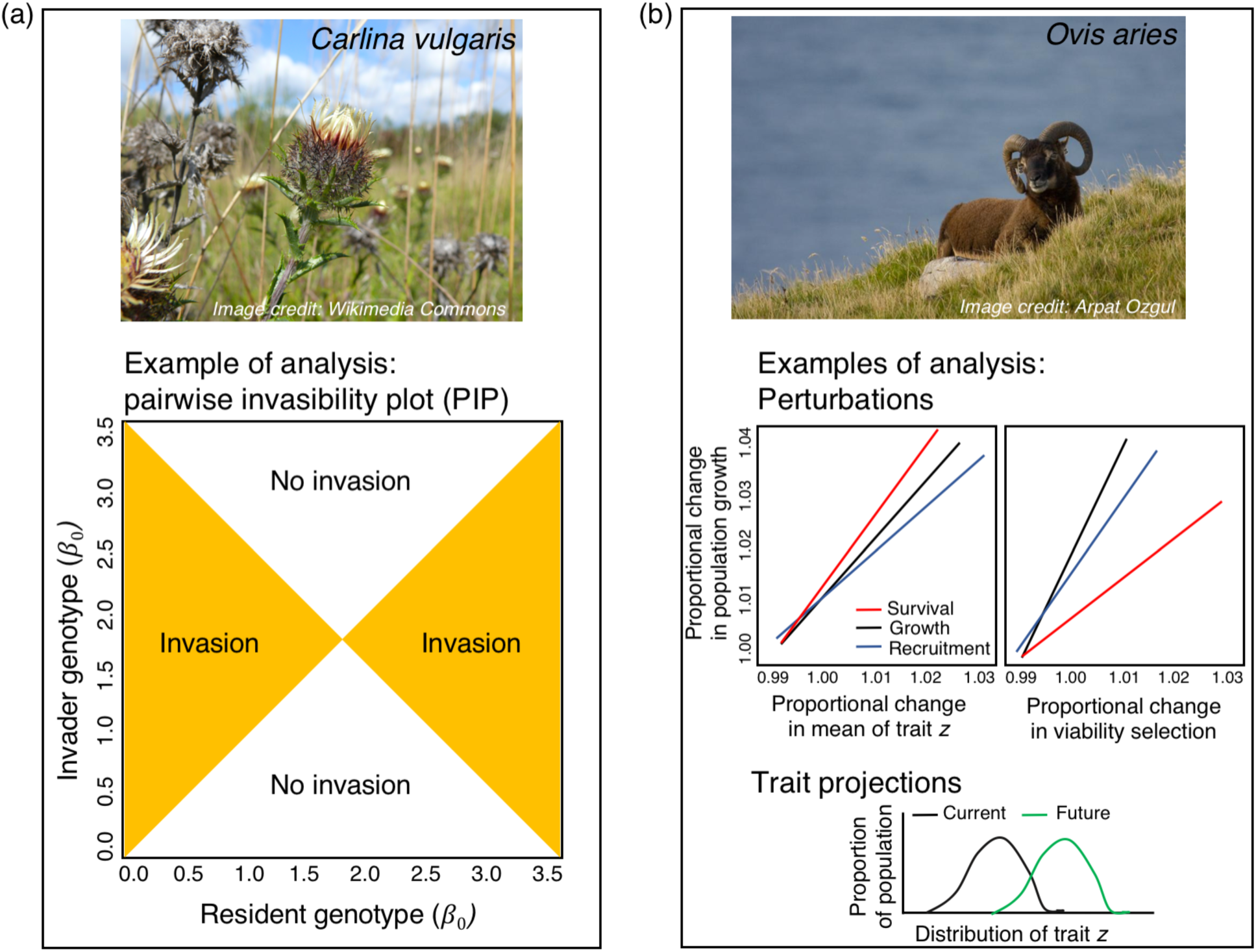
Examples of eco-evolutionary analyses using integral projection models (IPMs). For many plant species (a), e.g., the monocarpic thistle *Carlina vulgaris*, analyses focus on finding evolutionary stable strategies, *i.e*., values of a parameter (*β*_*0*_) in the demographic-rate regressions included in IPMs (eqn. 1), which cannot be invaded. Other eco-evolutionary IPMs (b), used largely in animal (e.g., the Soay sheep *Ovis aries*) population studies, track distributions of phenotypic traits (*z*) through time and use perturbation analyses to disentangle contributions of ecological and evolutionary processes to population dynamics.

IPMs incorporating ESS typically assume that one parameter in a model describing a demographic rate or transition, for instance *β*_*0*_ in eqn. 1, represents the genotype of an individual (Fig. 2a). The observable phenotype corresponding to a genotype is then a measure of the continuous trait *z* (Rees & Ellner, 2016). Both trait (*z*) and genotype (*x*) can be integrated into IPMs by expanding eqn. 3, assuming that offspring perfectly inherit their parent’s genotype:

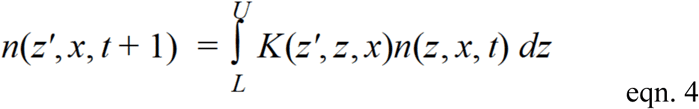

If inheritance is not complete, an inheritance kernel *M(x|x*^*P*^), describing the distribution of offspring of genotype *x*, given their parent (*P*) had the same genotype, must be integrated into the kernel *K* (Fig. 1).

### Eco-evolutionary dynamics

The focus on evolutionary endpoints does not inform about population responses to concurrent changes in phenotypic traits and ecological processes under global environmental change. For global-change research, such concurrent eco-evolutionary dynamics are of particular importance as an increasing number of studies have shown that evolutionary change can occur rapidly due to changes in selection pressure (Jump & Penuelas, 2005; Hoffmann & Sgrò, 2011; Schoener, 2011; Bonnet *et al.*, 2017). For instance, in the long-term Park Grass experiments at Rothamsted, experimental addition of soil minerals has resulted in adaptations of the perennial grass *Anthoxanthum odoratum* to liming, even when gene flow across treatments occurred (Snaydon & Davies, 1982; Silvertown *et al.*, 2005). Similarly, Ezard and colleagues (2009) have shown that variability in birth mass (phenotype) contributed strongly to the observed variation on the population growth rate of five ungulate species experiencing various levels of environmental variation. Therefore, in order to assess the mechanisms underlying life-history and population responses to global environmental change, phenotypic, and ideally explicitly genetic, changes through time must be tracked (Ozgul *et al.*, 2009; Coulson *et al.*, 2010, 2011; Smallegange & Coulson, 2013; Vindenes *et al.*, 2014).

Studies investigating dynamic trait distributions are mostly performed in animal populations (Vindenes & Langangen, 2015). Here, body size or mass, and not so much regression parameters, are seen as the individual characters of interest to answer joint evolutionary and demographic questions (Fig. 2b; Coulson *et al.*, 2010). Leaning on quantitative genetic approaches, IPMs can be used to infer traits and evolutionary metrics such as maternal effects, selection differentials, and heritability (Fig. 1b). For example, the Price equation (Price, 1970) has been integrated into the IPM framework in numerous studies to decompose changes in the mean and variance of a phenotypic trait into contributions from viability, development, and reproduction selection (Fig. 2b; Coulson & Tuljapurkar, 2008; Coulson *et al.*, 2010; Ozgul *et al.*, 2010). Such decomposition of contributions is typically accomplished with perturbation analyses of the parameters in demographic-rate functions (Fig. 2b).

### Measuring inheritance

Eco-evolutionary IPMs rely on explicit individual relatedness assessments using either observations or pedigree analyses (e.g., Leclaire *et al.*, 2013; Bonnet *et al.*, 2017). This allows for a straightforward investigation of the inheritance structure among traits. However, with some exceptions (Hanski & Saccheri, 2006; Santure *et al.*, 2013), we know very little about the individual relatedness and genetics underlying trait dynamics in plants. This is arguably in part due to the fact that mating and tracking of offspring are more difficult (Nyquist & Baker, 1991). However, as recent advances in plant studies show, there is in principle no obstacle to achieving more complex heritability estimates. Pedigree analyses and DNA markers to estimate genetic components of trait variation are increasingly common (Etterson & Shaw, 2001; Holland *et al.*, 2003; Jump *et al.*, 2008). For instance, Lloret & García (2016) detail genotyping procedures for kinship analyses among individuals of the small tree *Juniperus phoenicea*. As the following sections elaborate, the results of such efforts to genotype and relate individuals would be of critical importance for population models that are able to answer important questions of global-change research.

## Future applications of eco-evolutionary dynamics in plant demography

### Eco-evolutionary dynamics for conservation management under global change

Changes in the genetic and phenotypic makeup of a population that enhance population fitness, *i.e.*, adaptive plasticity, ultimately determine whether populations persist under environmental change (Paniw *et al.*, in review; Coulson *et al.*, 2017). For example, Coulson and collaborators (2011) assessed simultaneous changes in genetic traits (fur-color genotypes), body mass (phenotype), and demographic rates in wolves under environmental changes. The authors found that changes in environmental averages, but not environmental variation, can result in significant changes in eco-evolutionary dynamics in the wolf population. Such relative insensitivity to environmental variation may be related to the life history of wolves which are long-lived species with complex demography and are therefore expected to be buffered from increases in environmental variability (Morris *et al.*, 2008; Lenzi *et al*., in review). Other life histories, such as the Kalahari meerkats, which are relatively short-lived and inhabit unpredictably environments, exhibit strong changes in phenotypic trait distribution and population dynamics under increasing climate variability (Paniw *et al*., in review). Such trait-based life-history analyses are becoming increasingly relevant to unraveling the mechanisms of future population persistence (Paniw *et al*., 2017a; Paniw *et al*., 2018).

As in animals, global environmental change is affecting numerous plant populations (Thuiller *et al.*, 2005; Kelly & Goulden, 2008; Maestre *et al.*, 2012), and in order to persist, plants must either adapt or migrate (Aitken *et al.*, 2008). Increasingly, numerous mechanisms, including adaptation and phenotypic plasticity, that stabilize plant populations and communities, instead of driving them to extinction, are recognized (Nicotra *et al.*, 2010; Lloret *et al.*, 2012). For example, using population genetic analyses, Jump *et al.* (2008) discovered that rapid genetic adaptation to increased drought occurs in the Mediterranean shrub *Fumana thymifolia*, potentially increasing the probability of population persistence. Rapid evolutionary responses to environmental change may be particularly important for conservation management as they can induce genetic rescue, where adaptations to novel conditions may buffer population collapse (reviewed in Ellner, 2013).

Just as in animal populations, understanding trait dynamics in plants will be key to understanding eco-evolutionary dynamics. For instance, plant size has been shown to be a key determinant of vital rates and population dynamics across numerous ecosystems (Salguero-Gómez *et al.*, 2012; Paniw *et al.*, 2017a; Quintana-Ascencio *et al.*, 2018). Studies have shown potential shifts in size distribution due to shifts in selection pressures under environmental change with potentially significant consequences for conservation management (Paniw *et al.*, 2017a). The only way to understand the consequences of eco-evolutionary dynamics is however to incorporate trait dynamics into population-level processes (Reed *et al.*, 2013). It is surprising therefore that few IPMs are used for conservation management of plants, and, to my knowledge, models incorporating trait dynamics into eco-evolutionary feedbacks do not exist altogether.

To begin with, complex genetic models are not required to use plant traits, such as size, to inform on potential conservation management. Changes in the distribution of phenotypic traits (Fig. 2b) can used to improve predictions of early-warning signals of population collapse (Clements & Ozgul, 2016; Clements *et al.*, 2017). Whether changes in traits can predict population collapse in plants is little explored and provides an interesting novel avenue of research (Clements & Ozgul, 2018). Ultimately however, trait inheritance must be quantified to assess eco-evolutionary dynamics. Once this step is accomplished, numerous new avenues of research using eco-evolutionary IPMs are possible.

### Density-dependent feedbacks

Density feedbacks are important to accurately describe variation in demographic processes and population dynamics (Ellner *et al.*, 2016). In terms of eco-evolutionary feedbacks, it has been recognized that the strength of density dependence can strongly mediate the effect of selection pressures on phenotypes (Lowe *et al.*, 2017). For example, Reed and coauthors (2013) showed that a mismatch in phenology between food availability and egg laying dates in great tits (*Parus major*) creates a directional selection pressure towards earlier reproduction. However, higher chick mortality is compensated for by increased survival of the remaining chicks, which are released from negative density-dependent feedbacks. Such compensatory effects likely play an important role in nature but require integrating evolutionary dynamics into population models (Lowe *et al.*, 2017). Many studies in plant evolutionary demography looking at ESS have acknowledged that life history evolution is only poorly predicted when density feedbacks are omitted (Metcalf *et al.*, 2008; Gremer & Venable, 2014). Changes in population density or density of distinct life-cycle stages can also change the distribution of a density-dependent quantitative trait, thereby altering the population growth rate (e.g., Sæther & Engen, 2015; Travis *et al.*, 2015). However, density dependence is difficult to quantify in natural populations, and IPMs on plant populations typically omit density feedbacks. The consequences of such omission require further research.

### Dispersal

Another key aspect of population dynamics that is often ignored in plant and animal IPMs alike is dispersal. Migration patterns have been shown to strongly affect phenotypic traits and vital rates and may play a key role in evolutionary dynamics (e.g., Maag *et al.*, 2018). It is increasingly recognized that dispersal is not random but that certain genotypes or phenotypes are more likely to disperse than other (Edelaar & Bolnick, 2012; Ozgul *et al.*, 2014; Deere *et al*., 2017). For example, dispersers in some butterfly species consist of genotypes with greater flying capacity (Hanski, 2011). In desert plants, seed genotypes that promote short dispersal may create selection pressures towards more facilitative interactions under increasing environmental harshness (Kéfi *et al.*, 2008). However, spatially explicit IPMs are scarce (Merow *et al*., 2014b) and ones incorporating evolutionary mechanisms do not exist to my knowledge.

### Individual heterogeneity

The focus on one phenotypic trait, typically body mass, in eco-evolutionary IPMs received some critique as the growth (or trait transition) functions of IPMs (*G* in Fig. 1) do not track individual phenotypes through life and therefore do not account for the accumulation of phenotypic differences between individuals throughout life (Chevin, 2015; Janeiro *et al.*, 2017). If inheritance is then measured as a correlation between offspring and parent body mass, any genetic effect is confounded with ontogeny because older and heavier parents will have heavier offspring than younger ones, regardless of their own mass at birth (Vindenes & Langangen, 2015). This can lead to inaccurate estimates of inheritance, particularly for organisms with indeterminate growth (de Valpine *et al.*, 2014). However, IPM theory and applications are being continuously expanded, for example to embed quantitative genetic models of inheritance, which allow for better approximations of micro-evolutionary dynamics (Childs *et al.*, 2016; Coulson *et al.*, 2017).

One evident expansion of the eco-evolutionary IPM framework is to incorporate age structure into IPMs (Coulson & Tuljapurkar, 2008; Coulson *et al.*, 2010). Here, heritability of a trait such as size distribution at birth must be estimated for each age (Chevin, 2015); but even when this is done, explicit differences among individuals are not considered (Vindenes & Langangen, 2015). A more comprehensive approach is therefore to include static traits representing individual heterogeneity into IPMs (Fig. 1b) (Coulson *et al.*, 2011; Vindenes *et al.*, 2012, 2016; Vindenes & Langangen, 2015). Such traits do not change throughout the life of an individual, and may include traits such as size at birth, sex, or genotype (Vindenes & Langangen, 2015). This idea makes two relatively simple expansions of the IPM formulation in Fig. 1b: (a) creating a joint distribution of the recruit size distribution (*D*) and inheritance (*M*) functions and (b) incorporating a Dirac delta function into the *P* kernel: δ(*x*′ − *x*). This latter function allows to keep the static trait *x* constant for an individual during its lifetime (Vindenes & Langangen, 2015).

### Extrapolation of experimental approaches to population dynamics

In general, evolutionary responses to global change will be difficult to estimate from observational field studies alone because one must quantify not only all potential evolutionary reaction norms to a driver, e.g., temperature, but also the responses in continuous genetic and phenotypic traits to such drivers (Coulson *et al.*, 2011; Ellner *et al.*, 2016). However, it is possible to investigate such relationships in controlled laboratory or field experiments (Benton *et al.*, 2007; Collins, 2013), although it is difficult to detect the speed of eco-evolutionary feedbacks even in experimental studies (Ellner, 2013). Indeed, numerous studies have experimentally tested adaptations of plants to rapid environmental change (Jump & Penuelas, 2005; Jump *et al.*, 2008); however, the incorporation of experimental data into population-level models, including IPMs, is scarce (but see Olsen *et al.*, 2016). This is surprising and poses potential challenges to global change research as it has been increasingly recognized that negative effects of the environmental changes on vital rates can be buffered through complex tradeoffs and feedbacks, e.g., density dependence as described above or demographic compensation (Doak & Morris, 2010; Lloret *et al.*, 2012; Villellas *et al.*, 2015).

### Investigating selection pressures

Before detailed heritability information is collected, IPMs prove an ideal tool to investigate selection pressures on traits (Rees & Ellner, 2009). Seminal work by Tuljapurkar and colleagues (Tuljapurkar *et al.*, 2003; Haridas & Tuljapurkar, 2005; Horvitz *et al.*, 2010) has laid the ground for the assessment of selection pressures in structured population models in time-varying environments. This assessment is based on prospective perturbation analyses of demographic rates (*i.e.*, absolute or proportional sensitivities of the stochastic growth rate to these rates), and has been successfully applied, for example in the understory shrub *Ardisia escallonioides* or red deer *Cervus elaphus*, to test for direction selection on average vital rates and selection on vital-rate plasticity (Tuljapurkar *et al.*, 2003; Haridas *et al.*, 2009). Results from empirical analyses of selection pressures point to a tradeoff in evolving a sensitivity to changes in average vital rates (often found in long-live species) and their plasticity (more common in short-lived or disturbance-adapted species) (Haridas & Tuljapurkar, 2005; Morris *et al.*, 2008). The framework has been expanded to IPMs, which allows for straightforward trait-based assessments of selection pressures (Rees & Ellner, 2009). Empirical analyses have shown for example that plasticity in size is not favored in the thistle *Carlina vulgaris* (Rees & Ellner, 2009), while more variable seedling size may be beneficial for the carnivorous Mediterranean subshrub *Drosophyllum lusitanicum* under certain disturbance regimes (Paniw *et al.*, 2017a). Whether selection pressures translate into genetic changes can then be tested in experimental settings, as has, for instance, been shown for seed-bank dynamics in *D. lusitanicum* (Gómez-González *et al.*, 2018). Investigating selection pressures before or concurrent with detailed genetic analyses is therefore highly relevant to global-change research (Vindenes *et al.*, 2014) and should find more empirical applications beyond the studies presented here.

## Conclusions

In virtually all ecosystems, anthropogenically driven changes to the environment are threatening natural populations. Responses to these threats are complex and may include adaptations to novel conditions. To project future population dynamics, global-change ecological research must therefore use a flexible framework to disentangle the contributions from ecological processes (i.e., demographic responses to environmental change) and adaptive mechanisms (changes in genetic and phenotypic trait distributions) to these dynamics. This article detailed how integral projection models (IPMs) can function as such as versatile framework. Importantly, plant population ecologists can use much of the IPM methodology developed in animal population models to answer key questions in how plant populations may respond to global change.

## Acknowledgements

I thank C. García for the constructive feedback.

